# Temporal and spatial factors that influence magnetotaxis in *C. elegans*

**DOI:** 10.1101/252700

**Authors:** A.G. Vidal-Gadea, C.S. Caldart, C. Bainbridge, B.L. Clites, B. Palacios, L.A. Bakhtiari, V.D. Gordon, D.A. Golombek, J.T. Pierce

## Abstract

Many animals can orient using the earth’s magnetic field. In a recent study, we performed three distinct behavioral assays providing evidence that the nematode *Caenorhabditis elegans* orients to earth-strength magnetic fields (Vidal-Gadea et al., 2015). In addition to these behavioral assays, we found that magnetic orientation in *C. elegans* depends on the AFD sensory neurons and conducted subsequent physiological experiments showing that AFD neurons respond to earth-strength magnetic fields. A new behavioral study by Landler et al. (2017) suggested that *C. elegans* does not orient to magnetic fields and raises issues that cast doubt on our study. Here we reanalyze Lander et al.’s data to show how they appear to have missed observing positive results, and we highlight differences in experimental methods and interpretations that may explain our different results and conclusions. Moreover, we present new data from our labs together with replication by an independent lab to show how temporal and spatial factors influence the unique spatiotemporal trajectory that worms make during magnetotaxis. Together, these findings provide guidance on how to achieve robust magnetotaxis and reinforce our original finding that *C. elegans* is a suitable model system to study magnetoreception.

## INTRODUCTION

Most research on the neuronal basis for magnetosensation has focused on animals that migrate long distances by in part using the Earth’s magnetic field as a cue (Johnsen and Lohmann, 2005; Guerra et al., 2014). Although migrations by birds, butterflies, and turtles are magnificent in their own right, elucidating the cellular and molecular bases for magnetosensation is challenging in these complex animals.

Favoring a simpler animal, we recently asked whether the nematode *Caenorhabditis elegans* was capable of magnetic orientation (Vidal-Gadea et al., 2015). *C. elegans has proven historically important for the discovery of molecules used to sense odors, mechanical f*orce, osmolarity, and humidity (Sengupta et al. 1996; O’Hagan et al., 2005; Colbert et al., 1997; Russell et al., 2014). Notably, each of these molecules share conserved functions in higher animals (Tobin & Bargmann, 2004; Arnadóttir & Chalfie, 2010; Filingeri, 2015). If *C. elegans* displayed magnetoreception, potentially conserved molecular bases for this sensory modality may be studied using similar approaches. Using three distinct behavioral assays, we discovered that this tiny worm could orient its movement to artificial magnets or to the earth’s magnetic field (Vidal-Gadea et al., 2015).

By leveraging genetic tools specific to *C. elegans*, we next determined that the AFD sensory neurons were required for magnetic orientation and began to uncover signal transduction components required for magnetosensation (Vidal-Gadea et al., 2015). We also found that similar to magnetotactic bacteria, *wild C. elegans* strains isolated from around the world oriented to magnetic fields in a manner reflecting the magnitude and direction of the geomagnetic field at their point of isolation (Blakemore, 1975; Vidal-Gadea et al., 2015). Our behavioral and physiological results provide strong evidence that *C. elegans* senses and orients to magnetic fields.

Recently, Landler et al. (2017) performed additional sets of behavioral experiments to confirm whether *C. elegans* orients to magnetic fields. They reported negative results for all three experiments and conclude that *C. elegans* is not a suitable model system to study the molecular basis for magnetoreception. On first inspection, the experiments done by Landler et al. (2017) resemble those from our study with additional levels of control. We discuss subtle but important differences in experimental methods, controls, and execution that might have contributed to their negative results. We also discuss how behavioral results obtained by independent labs are consistent with our original findings. To help groups get started with *C. elegans* magnetotaxis, we present a new spatiotemporal analysis of worms orienting to magnetic fields along with suggestions on how to simplify our assays and control factors to ensure more robust results.

Lastly, Landler et al. (2017) discuss conceptual issues with our findings and interpretations. First, they suggest that directional information is absent in the magnetic field used in our magnetotaxis assay. Second, they suggest that a tentative explanatory hypothesis that we put forward– that *C. elegans* strains isolated from different locations on the globe may migrate at a specific angle to the magnetic field, perhaps as a way to orient optimally up or downwards when burrowing - is infeasible. We address these two issues by showing data from experiments demonstrating that the magnetic field does provide directional information in our magnetotaxis assay. Furthermore, thanks to this challenge, and the new experiments it prompted, we are now able to reconcile the results from our uniform and radial magnetic field experiments. We finish by identifying plausible mechanisms for how worms may use the directional information provided by a magnetic field to migrate along a specific vector.

Overall, the new data and updated interpretations offered in this response will likely help readers understand the discrepancy between the results reported in Landler et al. (2017) and those reported by our three groups. With this additional information, we hope that more researchers will be drawn to study the cellular molecular basis for magnetoreception using *C. elegans*.

## RESULTS AND DISCUSSION

This response is divided into four sections. We begin by summarizing of our previous results. The second section will discuss overt and potential differences between the assays conducted by Vidal-Gadea et al. (2015) and Landler et al. (2017). Third, we provide additional experiments and results from an independent lab confirming our original findings. In the last section, we discuss conceptual issues related to trajectories that worms make during magnetic orientation.

### Summary of previous results

Our previous paper included a large number of experiments and controls ranging from cellular calcium recordings, neuronal ablations, and cell-specific gene rescues, mutant analyses, and behavioral studies. The present challenge restricts itself to our behavioral experiments. We developed and performed <u>three distinct behavioral assays</u> to test whether *C. elegans* orients to magnetic fields (Vidal-Gadea et al., 2015) summarized below.

#### Burrowing assay

We tested whether British (N2) strain worms displayed a preference for burrowing up or down when injected into an agar-filled cylinder. We found that worms preferred to migrate *down* when starved (<u>which we defined as 30 or more minutes away from food</u>), and *up* when well-fed (<u>which we defined as less than 30 minutes away from food</u>). To test whether vertical bias was controlled by gravity or the magnetic field, we experimentally inverted an earth-strength magnetic field and found that the directions of migration were reversed. In Australia, the earth’s magnetic field has a polarity that is opposite that in Britain; we found that worms isolated from Australia migrated in the opposite direction to British worms. These results are consistent with the idea that the magnetic field, not gravity, provides the major cue for burrowing direction.

#### Horizontal plate assay

We tested how worms migrate in a uniform magnetic field generated by a magnetic coil system across a 10-cm diameter assay plate. We found that British worms moved randomly when the earth’s magnetic field was cancelled by the coil system, but migrated at a single specific angle with respect to magnetic north in the presence of an earth-strength field directed across horizontally positioned assay plate. Analogous to our Burrowing Assay results, the angle that worms migrated with respect to the field depended on the satiation state of the worms as well as their global site of isolation. When starved for 30 minutes, British worms migrated 180° away from the direction they migrate when well-fed. Australian worms migrated with a pattern opposite to British worms. Hawaiian worms migrated at a shallower angle, consistent with a shallower angle of the earth’s magnetic field with respect to the local surface of the Earth. Considering the horizontal position of the assay plate and the opposite migration preferences of worms from northern and southern hemispheres, these results support the idea that *C. elegans* detect and orients to magnetic field in accordance with its global site of isolation.

#### Magnetotaxis assay

We tested whether British worms migrated towards or away from an artificial magnet placed north-side up beneath one side of a 10-cm diameter assay plate. We found that well-fed worms migrated towards the magnet and randomly in the absence of a magnet. Genetic ablation of AFD sensory neurons rendered worms unable to orient to the magnetic field. Additional analysis of mutant and cell-specific rescue strains revealed that the cGMP-gated cation channel TAX-4 was necessary and sufficient in AFD neurons for magnetic orientation because selective rescue of TAX-4 in AFD in a *tax-4* null mutant background rescued magnetic orientation. Our finding that all strains except for those with defects in AFD neurons performed magnetotaxis suggests that this behavior reflects the ability to orient to the field rather than an artifact of the magnet non-specifically influencing the movement of worms.

### Overt differences in experimental methods

Recently, Landler et al. (2017) attempted to replicate our results with British worms by modifying our three experiments to include worthwhile control measures and analysis that differed slightly from our original study. Unfortunately, it appears many of these experiments deviated from our described methods. For each experiment, they found negative results concluding that *C. elegans* may not orient to magnetic fields. Below, we discuss differences in experimental methods, analysis, and interpretation that may explain their failure to replicate our results.

#### Animal satiation states

The major difference between our methods was in the duration of the assays, and consequent changes in animal satiation. In Vidal-Gadea et al. (2015) we experimentally determined and reported that 30 minutes away from food was sufficient to flip the magnetotaxis behavior of the worms from positive to negative. This was initially unexpected because *C. elegans* does not flip its orientation preference after 30 minutes away from food for other orientation behaviors including chemotaxis to benzaldehyde. We therefore described this time as sufficient to induce the ‘starved’ state in animals, and went on to perform several experiments with worms in the ‘fed’ or ‘starved’ states. For ‘starved’ assays, we ensured that worms were away from food for 30 minutes prior to starting an experiment. From reading Landler et al. (2017) it is now clear that we did not explicitly mention that this definition of starvation implied that for worms to be tested in the ‘fed’ state, animals would need to complete their assay within 30 minutes.

We found that with practice by controlling the set up and number of worms we could run our behavioral assays to near completion within this 30-minute time window and did not need to immediately tally immobilized animals. This allowed us to run many assays in parallel without having to stop to conduct the time-consuming tallying step for each pipette or plate before starting the next. Tallying animals in the magnetotaxis assay, however, was a much simpler (and faster) procedure, which we could do at the 30 minute mark. Therefore, it is important to note that we ran assays so that worms migrated to a particular direction within 30 minutes.

It is clear from Landler et al. (2017) that they decided to modify our magnet assay to last <u>60 minutes</u> rather than 30 precisely because they continued to see moving animals all the way until this time point (see their Methods). Unfortunately, this also implies that many worms participating in the assay (which they described was a sufficiently large number to make them deviate from our protocol) would have transitioned to the ‘starved’ state. By our described definition of ‘fed’ and ‘starved’ (also adopted by Landler et al.) they report
testing animals under both ‘fed’ (first half of the assay), and ‘starved’ (second half of the assay) conditions. This issue may have been obviated when Lander et al tested prestarved worms (their Fig 3B); however, no-magnet controls and horizontal-oriented tube controls for these assays were not reported (see below). We believe this singular, and crucial, difference might explain the different results by Landler et al. Intrigued by this possibility, we conducted a new set of experiments where we film animals orienting in a double-wrapped magnetic coil system assay over time and report their shift in both the direction and consistency of the response over a 90-minute window in line with our observations, and those reported by Landler et al. (see below).

**Figure 1.**
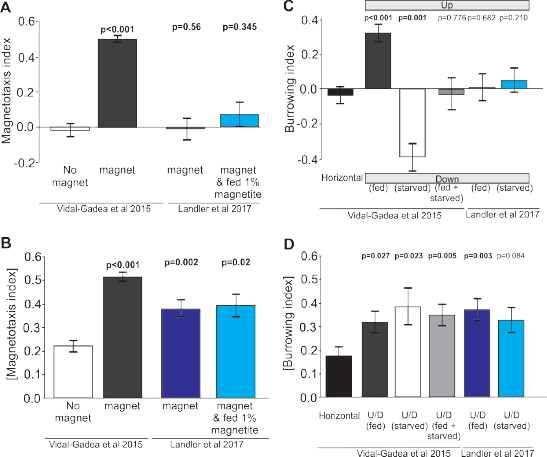
Reanalysis of data suggests that positive results may have been masked in Landler et al. (2017) by testing worms in both fed and starved states and omitting controls. **A)** Comparison of magnetotaxis data reported by Vidal-Gadea et al (2015) and Landler et al (2017) obtained by measuring their plots. Because Lander et al omitted nomagnet control data, we used no-magnet control data from Vidal-Gadea et al (2015). We found that Landler et al (2017) worms fed OP50 bacteria (or OP50 plus 1% magnetite) show no significant orientation versus no-magnet control worms. **B)** Under the hypothesis that Landler et al might have combined fed and starved worms because their assays were run for twice as long, we used the absolute value of the magnetotaxis index to reveal evidence that worms display a biased migration in the presence of a magnetic field (irrespective of the towards or away sign of their migration). We found that both magnet treatments in Landler et al. (2017) resulted in significantly biased migration when compared with no-magnet controls. **C)** We also analyzed burrowing data from Landler et al. (2017) and used our horizontal controls because they were omitted in Lander et al., (2017). We demonstrate that combining data from fed and starved worm abolished significant burrowing indexes that were otherwise observed from each of these populations. Similarly, comparison of Landler et al (2017) burrowing indexes to our horizontal controls (N = 24) revealed no burrowing bias in their field up results for either fed or starved conditions. **D)** However, when we compared the absolute value of burrowing bias we found that our combined fed ± starved group, as well as Landler et al’s “fed” worms now showed significant bias when compared to horizontal controls. All tests based on Mann-Whitney Ranked Sum Tests.

**Figure 2.**
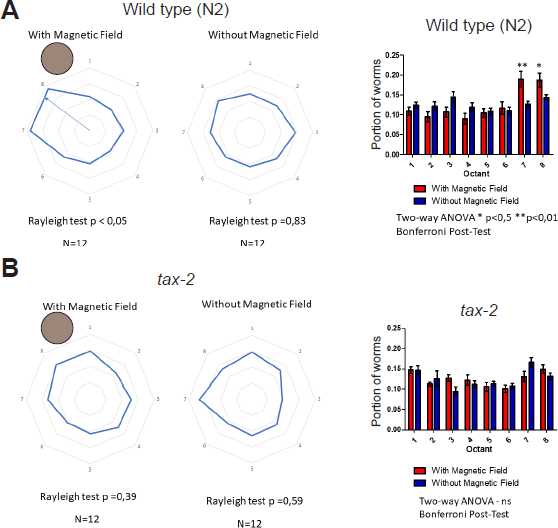
Independent evidence for magnetic orientation in *C. elegans*. Caldart and Golombek at the University of Quilmes, Argentina performed a modified version of the magnetotaxis assay described in Vidal-Gadea et al. (2015). Well-fed worms were placed at the center of a plate seeded with bacterial lawn with no sodium azide anesthetic. The fraction of worms found in each octant at 15 minutes is reported on the right. A strong neodymium magnet was placed adjacent north-side up. **A)** Wild-type N2 worms spent more time in octant to the left the magnet (red bars) compared to paired control assays where there was no magnet (blue bars). **B)** By contrast, *tax-2* mutant worms showed no preference for any octant in magnet versus no magnet conditions. N= 12 assays for each condition. Bars represent s.e.m.

**Figure 3.**
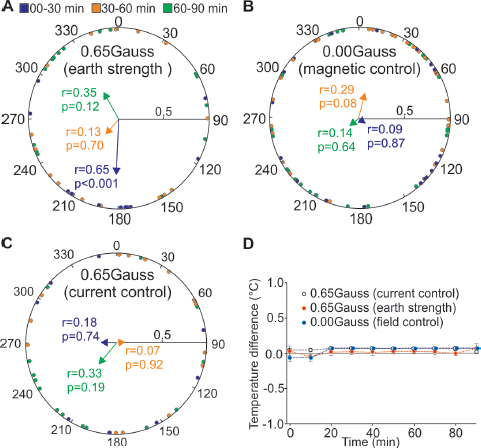
Time dependence of magnetic orientation in *C. elegans*. We tracked the centroid of freely moving worms in our double-wrapped magnetic coil system. **A)**Well-fed worms displayed robust magnetic orientation when placed in a 0.65-Gauss magnetic field (1xearth) for about 30 minutes (183°, r = 0.65, p < 0.001). After this time point their orientation became markedly random (r = 13, p = 0.7), only to consolidate again into the opposite direction (341°) after 60 minutes (r = 0.35, p = 0.12). **B)** Exposing the worms to a 0.00 Gauss field inside the coil system (by cancelling out earth’s field) resulted in random migrations. **C)** Similarly, powering the coil system under similar conditions as in **A** but switching the coils into their anti-parallel arrangement used similar currents to generate two antiparallel magnetic fields that cancelled each other out (current control) and produced random migration in test animals. **D)** We measured temperature gradients within the coil system over the course of our 90-minute assays and found that temperature differentials between the center and edge of the plate were below the known thermal sensitivity or worms. Headings calculated using individual assay means (number of assays: 1xearth N = 7, magnetic control N = 10, current control N = 6). Animal headings were tested using Circular Toolbox for MATLAB. Rayleigh tests measuring circularity. Temperature measurements were obtained by using high sensitivity thermometers as previously described (Vidal-Gadea et al., 2015).

#### Temperature controls

The absence of temperature recordings makes it difficult to predict if this important variable was sufficiently controlled by Landler at al. We note that in their Fig 3D the resulting vector they obtained under test conditions is consistently twice that of their control conditions. This difference may have reached significance had they not terminated their assay after 30 minutes and/or increased sample size. Thermal gradients within the test chamber may have differed in control versus test conditions.

### Additional differences in experimental methods

We next describe additional differences between our experiments that might have further contributed to their observations.

#### Burrowing assay

Landler et al. (2017) assayed whether N2 worms injected into agar-filled cylinders burrowed up or down. They report no bias for burrowing up or down, with or without an imposed inverted magnetic field.

First, regarding experimental order: before we attempted to manipulate the magnetic field surrounding the pipettes we performed our experiments in electrically shielded pipettes aligned vertically in the lab, and in the absence of artificial electric and magnetic fields. It was only after obtaining positive results in these conditions that we proceeded to experiments using a magnetic coil system.

Second, regarding the assay vehicle: we used plastic (5 mL) serological pipettes and Parafilm to seal the pipette perforations because it prevents desiccation but allows free gas diffusion. We are uncertain regarding their assay tube material, but understand that they used plastic caps which presumably prevented both water and gas exchange. Depending on the number of animals, sealing the pipettes might contribute to the generation of CO_2_ and O_2_ micro-gradients that might affect animal behavior.

Third, as correctly pointed out in Landler et al. (2017), our original magnetic coil system differs from theirs in not being double wrapped. Nevertheless, we recently used a new double-wrapped coil system in the Vidal-Gadea lab at Illinois State University to successfully replicate our earlier findings (see below and Fig 3). Another difference in coil systems arises from their geometry. In our 2015 study, we used three sets of four-coil systems described by Merritt (Merritt et al., 1983) which was reported to produce more stable test fields for larger samples (e.g. 50 vs 20 cm^3^) than with Helmholtz loops (Kirschvink, 1992; Magdaleno-Adame et al., 2010) used by Landler et al. (2017). Before we conducted our burrowing assays inside our magnetic coil system, we measured the field properties throughout the test volume to ensure that our system produced a homogeneous field. Lander et al. (2017) did not report such measurements of homogenity.

Fourth, Landler et al. (2017) suggest that unintended temperature gradients generated by our coil system may have resulted in our reported observations. However, aware of this possibility, we used high sensitivity thermometers (0.01°C) to quantify temperature changes (in both magnet, and magnetic coil system assays). A two-way ANOVA (N = 5, p = 0.123) showed no significant difference in temperature between cancelled field and one-earth field (Fig 2. Supplement 1 in Vidal-Gadea et al., 2015). As an extra precaution, we regularly rotated the orientation of our magnetic coil system to a random position before each assay, and used a small fan to circulate air through it and prevent temperature gradients. Landler et al. (2017) do not describe doing any such controls and they do not present associated measurements. However, from their discussion they seem to believe that the use of double wrapped coils prevents the generation of meaningful temperature gradients. This is incorrect; all current passing through a metal conductor generates heat. Therefore, under both their test and control conditions, heat would be generated by current passing through their system potentially building up with the small enclosed metal room used to shield worms from foreign magnetic fields. We welcome their additional control of a mu-metal shielded room as an improvement to minimize potential magnetic contamination, but caution that environmental parameters should be controlled further by positioning the coil system at random orientations within the room, and by empirically measuring and mitigating the thermal gradients that must necessarily develop as current powers any coil system as we did in our original study.

Fifth, our burrowing experiments tested in the natural earth field showed that worms migrated differentially based on their global site of origin and satiation state. This observation undermines the likelihood of temperature gradients, or magnetite contamination, being responsible for our results. Furthermore, our findings that worms lacking the transduction channel encoded by the *tax-4* gene, or by worms with genetically ablated AFD neurons, failed to burrow preferentially up or down in earth’s natural field strongly point to the involvement of these neurons and molecules in this behavior (Vidal-Gadea et al., 2015). None of these results are mentioned in Landler et al. (2017).

#### Horizontal plate assay

Landler et al. (2017) tested how worms migrate to the edge of a 10-cm diameter plate in a horizontal magnetic field where they were trapped by azide at the edge. Unlike our study, they found no significant degree of orientation in their migration.

In Landler et al. (2017), they point out that we treated each worm as an individual in our horizontal field assays when performing statistical analysis, which could cause a type 1 error if worms did not act independently. While we believe that the individual timing and trajectory of each worm makes them independent, which justified our choice of statistical analysis, we nevertheless re-analyzed our data averaging the mean heading of worms in each assay as they did. Similar to our previous report, we found that in all but one out of 28 assays, worms displayed a significant migratory preference. The average heading of the assays changed somewhat from the result reported in Vidal-Gadea et al. (2015), but not significantly so. Furthermore, we provide new experimental data following Landler et al.’s suggestion for analysis and show similar results (see below).

Landler et al. (2017) also noted that all worms in their magnetic field conditions are set by an experimenter not involved in the analysis. This blinding protocol was also the case for our original study, but not explicitly mentioned. Anecdotally, we expected British worms to migrate towards magnetic north, just like magnetotactic bacteria, but remained puzzled for months when our results unexpectedly showed them consistently migrating at a 132° angle with respect to magnetic north. This illustrates how our expectations did not affect our analysis or results. Indeed, we still do not know why worms prefer this particular angle, although we presented a parsimonious explanation that worms may choose a migratory direction based on the inclination of their native field.

#### Magnetotaxis assay

Landler et al. (2017) attempted to replicate our magnetotaxis results and reported that worms migrated randomly towards and away from the magnet.

In their manuscript, Landler et al. (2017) offered no actual replication for our magnetotaxis experiments, opting instead for modified protocols. They described using a brass coin as a control in their magnet assays. We similarly used a cylinder fashioned of aluminum foil and similar dimensions to the test magnet under the control area in our assays. Landler et al. chose to extend their assay time to one hour to include worms that may have switched to a starved state. As described above, it is therefore likely that by testing fed and starved animals alongside one another, they effectively combined and measured positive and negative magnetotaxis.

Landler et al. conducted an additional control in the form of a magnetic assay with worms fed 1% magnetite throughout their cultivation. They reported a barely significant (p < 0.04) improvement over chance performance data that were not plotted; however, their assay was still not significantly different from animals that migrated in the presence of a magnet but were not fed iron. Their observation that worms purposely contaminated with magnetite fail to migrate to a magnet as in our assays provides evidence that contamination with magnetite is likely not responsible for our observations. However, we take this conclusion with caution since their one hour assay might have masked potentially significant effects arising from their magnetite enrichment.

Magnetite is pervasive in the decaying fruit and plant matter where worms live. It is likely that worms in their natural habitat may have access to much greater amounts of iron than those provided for them in the lab via a diet of *E. coli* on an agar surface. If *C. elegans* does use magnetite to build its own magnetic field detector, it is possible that enrichment of the lab culture conditions may result in increased magnetic indices for adults, or even succeed in enabling larval stage worms to perform this behavior better than what we previously reported (Bainbridge et al., 2016). While we consider this experiment informative, we do not agree with Landler et al. (2017) in their suggestion that ingestion of magnetic particles might confer *C. elegans* (or other animal for that matter) the ability to migrate within magnetic fields. This would not explain how worms migrate at different prescribed angles in a satiety-dependent manner, or that this behavior can be systematically abolished and rescued by cellular and molecular tinkering. To our knowledge, ingestion of magnetite or iron is yet to be demonstrated <u>in any animal</u> to be sufficient to confer the ability to migrate towards and or away from magnets using their own power. Thus, although exogenous iron has been suggested to contaminate cells proposed to be magnetoreceptors (e.g. Edelman et al., 2015), in contrast to Lander et al.’s (2017) claim, there is no example of false positives associated with actual magnetoreceptive behavior or physiology displayed by any wild-type animal.

One important control provided in our manuscript but left out in Landler et al. (2017) was the response of worms in the absence of a magnet. In Figure 3F the authors display results for two assays in which worms are in the presence of a magnet, with the only difference being whether or not their diet was enriched with magnetite. No control assays where worms tested in the absence of a magnet are provided, and worms tested with a magnet are confusingly labelled “baseline”. We find the omission of this no-magnet control in particular, and the nomenclature used, concerning. Based on our experience, fed worms display magnetotaxis indices with high positive values, and starved worms display indices with high negative values. If Landler et al. inadvertently combined in effect starved and fed worms in their assays, then we expect them to observe a broad range of magnetotaxis indices centered at zero. These results would starkly differ from control assays with no magnet where indices would not stray far from zero (Vidal-Gadea et al., 2015). We suggest the authors plot this crucial control that they suggest that they conducted in their Results. If their no-magnet control data are significantly more narrowly distributed around zero than their test data, this would suggest that the broad distribution of indices obtained in their magnet assays is likely the result of combining fed and starved animals that are in fact orienting to the magnetic field.

To investigate this possibility that Landler et al. had obtained positive results that were masked by testing worms in both fed and starved states, we reanalyzed their data from Figure 3F. First, we plotted their original results in bar format alongside results from our original study including our no-magnet control (Figure 1A). Next, we plotted the absolute value of magnetotaxis indices (Figure 1B). With this alternative analysis we found that worms migrating in the presence of a magnet displayed a significant migratory bias when compared to worms migrating in the absence of a magnet (Figure 1B). Landler et al. (2017) reported that their magnetite-enriched worms did not migrate significantly better than worms that were not fed magnetite (p = 0.652). Our reanalysis confirms their finding, but also shows that their worms in both test conditions displayed a significant difference from our no-magnet control. We also reanalyzed burrowing data from Landler et al. (2017). Once again, we provided missing controls in the form of worms burrowing in horizontally oriented pipettes and compared these to data from both groups (Figure 1C&D). We found an identical pattern of results where the absolute value of the burrowing index of test group worms was significant different from the value for our horizontal control worms (Mann-Whitney Rank Sum Test, p = 0.003). Importantly, our reanalysis suggests that had no-magnet and horizontal controls been included and analyzed in this manner above, Landler et al. would have had to conclude that they obtained positive results for magnetic orientation in *C. elegans*.

### Important experimental variables for magnetic orientation in *C. elegans*

Below we list additional variables that may or may not differ between Landler et al. (2017) and our previous study. We have found that these variables matter for successful robust replication of magnetic orientation experiments with *C. elegans*.

#### Animal and environmental variables

Timing and satiation state are two of the most controllable variables affecting this assay. In Vidal-Gadea et al. (2015) we noted that crowding, ambient humidity, temperature, starvation, and contamination history could all sway the preference of a population from positive to negative magnetotaxis. This is perhaps not surprising given that the polymodal AFD neurons respond to temperature, humidity, and CO_2_ in a satiation-dependent manner (Mori, 1999; Bretscher et al., 2008; Russell et al., 2014). Indeed, we ceased to test animals during weather events (e.g. rain, high humidity, temperature fluctuations), and constructed an environmental box that maintains animals in constant temperature and humidity. Importantly, we have found that cultivating worms in an incubator (as was done in Landler et al. 2017) can interfere with the robustness of performance. Incubators cast strong magnetic fields that may affect worms during their course of development (Makinistian & Belyaev, 2018). Removing worms from an incubator with one temperature and magnetic conditions to different conditions in the assay room may shock worms.

Our experimental results show that it is crucial to know the state of the worms before and during assays. We hesitate to suggest which parameter may have caused trouble in Landler et al. (2017), although we venture to propose time/satiation state as an obvious starting place. We do know that once the correct physiological state has been produced in the animals, they will perform this behavior. Two lines of evidence substantiate this: **1)** special needs students, high school volunteers, and undergraduate students have all replicated our results in Texas and in Illinois when handed worms properly cultured; **2)** an independent group in Argentina recapitulated our results by conducting the experiments on a bacterial lawn (and thus avoiding the risk of on-assay starvation; see below).

Lastly, it is common practice to test multiple *C. elegans* strains that are freshly thawed from cryopreservation to verify the relation between genotype and phenotype. Although all results reported by Landler et al. (2017) used a single thaw of N2 strain obtained from the *Caenorhabditis* Genetic Center, it would add an additional level of control to test multiple biological samples. This would minimize the risk of testing a strain that developed spontaneous mutations that interfere with expected phenotypes.

### Challenge of assaying magnetic orientation in *C. elegans*

The field of magnetic orientation has just started in *C. elegans*, so it may not be surprising that parameters for robust replication of these assays are not yet optimized. By way of comparison, thermal orientation in *C. elegans* was first reported by Hedgecock and Russell (1975) and verified two decades later by the Mori lab (e.g. Mori and Ohshima, 1995). A decade further, experienced *C. elegans* labs (Goodman, Sengupta, and Samuel) subsequently replicated negative thermotaxis but failed to demonstrate positive thermotaxis (e.g. Clark et al., 2007). After a few years, these three labs working together with the Mori lab discovered that positive thermotaxis required a shallower temperature gradient (Goodman et al., 2014). Similarly, perhaps subtle aspects of the magnitude and shape of the magnetic field contribute to difficulty in Landler et al. (2017) in reproducing our experiments.

Despite these obstacles, we feel these challenges are worthy for researchers keen on seeking to study this fascinating sensory modality. For example, we are aware of at least two independent groups who devised different ways to overcome the fickle satiation and time dependence of the magnetotactic assays. Golombek and colleagues run their assays on plates covered in bacterial lawns, which removes the possibility of their worms starve. A second group interested in magnetotactic ability, rather than preference, calculates the absolute value of the magnetotactic index (Brenda Houck, Hendrix College, USA, personal communication). This simpler metric reflects the **magnetotactic ability** of a strain, as opposed to its *magnetotactic preference*. With this scoring system, magnetotaxis index values significantly greater than chance level without a magnet show an ability to detect and orient a magnetic field. This scoring was used in our reanalysis of Landler et al.’s data above (Fig 1). We consider both simplifications meaningful improvements over our initial assays and suggest their use to groups joining the field.

### New behavioral experiments support original study

#### Replication by independent labs

Since our initial description of this behavior in *C. elegans* (Vidal-Gadea et al. 2012a), we are aware of several groups joining the study of magnetic field detection using nematodes. While conducting our original study, Ilan et al. (2013) reported that parasitic nematodes migrated preferentially south over north when placed in a magnetic field. We recently became aware of a group at the University of Quilmes, Argentina who replicated our findings with minor modifications. Here we present their results, obtained in complete independence from the Vidal-Gadea and Pierce labs. Figure 2 shows Caldart and Golombek’s version of our assay. To avoid satiation shifts, this group conducted their assay in an agar plate seeded with *E. coli* bacteria and for only 15 minutes. Similarly to Vidal-Gadea et al. (2015), they observed a significant increase in worms congregating near the north-side facing up magnet (Fig 2A, Rayleigh clustering test, p < 0.05, n = 12) compared to controls (Rayleigh clustering test, p = 0.83, n = 12). Consistent with our original study that found that magnetic orientation required a cGMP-dependent signaling cascade including the cation channel TAX-4, Caldart and Golombek found that mutation of the TAX-4-obligate subunit TAX-2 abolished this migration (Fig 2B, p = 0.39, n = 12 in the presence of a magnet, p = 0.59, n = 12 when no magnet was present).

Landler et al. (2017) notes that a study by Njus et al. (2015) reported that worms failed to respond to magnetic fields. Njus et al. restricted their study to crawling velocity and omega bends and not orientation. Nevertheless, in Figure 6 of Njus et al. (2015), they show a 10±7% to 90±20% change omega bends when a 5-mT magnetic field was introduced or removed respectively. Such difference in turns would have a significant effect on course trajectory and orientation. Rather than comparing these paired measurements to each other, Njus et al. compared them to the number of worms turning in the absence of a magnetic stimuli for which they report an average of 70±50%. A 50% variability in omega bends is surprising and not consistent with previous reports in the literature (e.g. Vidal-Gadea et al., 2012b), or even with the variability they report for the rest of their data (22.5±8%, obtained by measuring and averaging standard deviations from test conditions reported by Njus et al., 2015: Fig.6). Not surprisingly, no test condition was significantly different from such a variable control. Therefore, in our view, the Njus et al. (2015) study appears to offer little evidence to counter the idea that *C. elegans* orients to magnetic fields.

Aware of the challenges presented by the study of magnetic orientation in *C. elegans*, the Vidal-Gadea and Pierce labs continue to study the time and environmental dependence of magnetic orientation in *C. elegans*. Below we provide example results demonstrating time-dependence of magnetic orientation from the Vidal-Gadea lab.

#### Magnetic field heading over time

To determine the effect of time on migratory angle we placed fed worms at the center of a Petri plate within our double-wrapped magnetic coil system. We used an electrically shielded USB camera to film worms migrating in a uniform magnetic field of 0.65 Gauss (1xearth) amplitude, and tracked the centroids of all worms participating in the assay. The average heading for all worms in the assay was noted during each 10-min interval over the course of 90 minutes. We hypothesized that if worms change their migratory angle as they become starved, then we should detect an increase in the variability of their migratory angle as time progresses and more worms enter a ‘starved’ state. We also predicted that over time, all worms would become starved and their migratory angle should tighten once again to a direction opposite their original preferred angle (as observed in Vidal-Gadea et al., 2015). Figure 3 shows the results for the headings of the animals over the course of a 90-min assay. Over the initial 30 minutes we measured worms mean heading of 183.2° (r = 0.65, p < 0.001). Between the 30 and 60-minute mark (time period tested by Landler et al. 2017) the mean direction of the population became scattered as many worms became starved (heading = 210.2°, r = 0.13, p = 0.70). Finally, beyond 60 minutes (60-90), most of the worms appeared starved and displayed a non-significant heading (341° avg, r = 0.35, p = 0.12). These results are both consistent with our previously reported findings (Vidal-Gadea et al., 2015) and with the reported measurements of Landler et al. (2017) considering the time window that they assayed worms.

Because electrical generation of magnetic fields in coils implies the unavoidable generation of heat, we used a fan to circulate air through our magnetic coil system and included two controls in our magnetic assays to control for magnetic field and temperature. We canceled out all magnetic fields inside our coil system to test worms in the absence of a magnetic field (magnetic control, Figure 2B). In addition, we controlled for heat by running the same current through the coil system in antiparallel configuration (temperature control, Figure 2C) thus generating a similar heat signature as that produced during our test. Importantly, we recorded and reported temperature changes in our system throughout each experiment. Temperature gradients inside our system were below the reported threshold for *C. elegans* sensitivity (Fig 3D).

### Conceptual issues regarding magnetic orientation in *C. elegans*

In addition to methodological issues, Landler et al. (2017) raise two conceptual issues regarding our original study that we address below.

### Magnetotaxis assay trajectories

How do worms move in our magnetotaxis assay? As described above, worms are placed in the center of an agar-filled Petri plate with a 0.29-T strength, 1.5-inch diameter, neodymium magnet placed north-side facing up 1-cm beneath the agar surface on one side of the plate (Fig 4 A-C). Azide is pipetted on magnet and control sides to immobilize worms that reach either location. Landler et al., (2017) suggest that the magnet in our magnetotaxis assay would generate field lines that pierce the surface of the agar assay plate only perpendicularly, as if worms were “on the magnetic north pole of the planet (with all directions being southerly), neither the polarity nor the inclination of the field can be employed by nematodes as a guidance cue” (see their Fig 4A). If so, then the magnetic field would provide no directional information. Below we explain how this viewpoint is incorrect.

**Figure 4.**
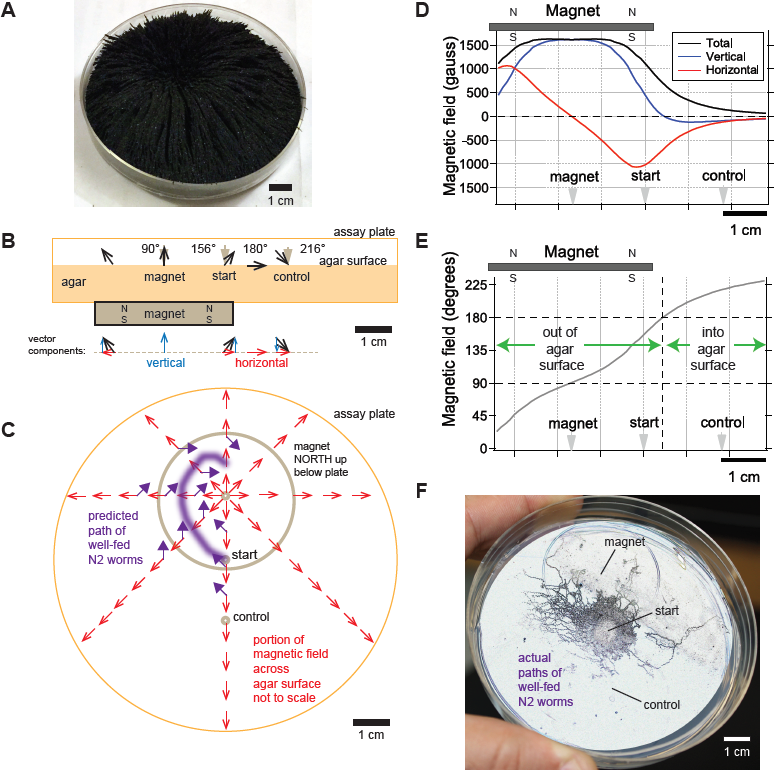
Directional information in magnetic field predicts *C. elegans* magnetotaxis trajectory. **A)** The direction of iron filings scattered across an assay plate reveals the general shape of magnetic field emanating from a 1.5” diameter magnet, north-facing up beneath the plate. **B)** Side view of magnetic field lines and their vertical and horizontal components across the surface of the agar-filled plate. Magnet and plate shown to scale. Field line strength not to scale. Gray arrowheads denote start location for worms and points where azide was spotted above the magnet and control goals. **C)** Top view of horizontal component of magnetic field (red arrows) across the surface of the agar-filled plate. Note that magnetic north points directly away from the center of the magnet everywhere on the plate. Wild-type N2 worms prefer to move at 132° to magnetic north, which predicts the trajectory (purple arrows and line). Field lines not to scale. **D)** Strength of the total magnetic field and its vertical and horizontal components across the agar surface. **E)** Inclination angle of the magnetic field across the agar surface. **F)** Majority of observed trajectories for N2 worms in the magnetotaxis assay arc left of magnet consistent with prediction.

To help visualize the magnetic field in our magnetotaxis assay, we scattered iron filings across the agar surface of the assay plate (Fig 4A). The filings stand straight up directly above the center of the magnet, indicating that the magnetic field is perpendicular to the agar surface here. Filings tilt at increasingly shallow angles as the distance from the magnet center increases until they become parallel with the plate surface. This simple experiment shows that magnetic field lines pierce the surface of the agar at a variety of angles in a radial pattern which could provide abundant directional information pointing away from the center of the magnet in our, and Landler et al.’s, experiments.

We next calculated the direction and magnitude of the magnetic field across the assay plate. This was done using the online calculator by K&J Magnetics, and validated experimentally with the aid of a magnetometer. We found that the magnetic field was strongest at the center of the magnet and began to strongly dissipate near the inner edge of magnet (Fig 4D). The horizontal component of the field that is parallel to the plate surface was zero at the magnet center and increased away from this point up until near the inner edge of the magnet (red line, Fig 4D). The sign of the horizontal component switched at the center of the magnet reflecting how the field lines point radially away from the magnet center. The angle of field penetration varied across the assay plate as expected from the iron filings (Fig 4A,B&E). The field pierces out of the agar surface at 90° only at the center of the magnet, and starts tilting until it reaches 180° about 21.5 mm from the center of the magnet (Fig 4B&E). Beyond this point, the magnetic field starts to tilt further, piercing into the surface of the agar. Therefore, the magnetic field in the magnetotaxis assay varies in polarity and inclination which we hypothesize worms may use as a cue to orient (Fig 4B&C). The magnetic field also varies in strength with levels far above the earth’s magnetic field. However, we found that the AFD magnetosensory neurons failed to generate larger responses when presented with larger than earth-strength fields (Vidal-Gadea et al., 2015).

Armed with an empirically validated model of the magnetic field in our plates, we can predict how worms move in the magnetotaxis assay. We previously found that in a uniform horizontal field, N2 worms preferred to migrate approximately 132° away from magnetic north when well fed (Vidal-Gadea et al., 2015). Given this preference, we expect that worms would make a leftward arc when viewing the assay plate from above (Fig 4C). This is because worms started at the center of the assay plate would consistently bear 132° away from the horizontal field lines (purple arrows, Fig 4C). To test this prediction, we retrieved photographs of assay plates from our original 2015 study. We found that a significant portion of well-fed worms migrate towards the magnet along the left side of the plates (Fig 4F). This unusually asymmetric arced trajectory contrasts greatly from the typical symmetric trajectory that worms make when migrating to the peak of an attractant chemical gradient during chemotaxis (e.g. Pierce-Shimomura et al., 1999).

To quantitatively test the prediction that worms would make left trajectories, we performed a modified version of the magnetotaxis assay using six symmetrical spots of azide to immobilize worms (spots A-F, Fig 5A). As before, the magnet faced north upward so that the horizontal component of the magnetic field pointed away from the magnet. A population of N2 worms was released at the center of the 10-cm plate and allowed to crawl freely for 30 minutes. The number of immobilized worms were tallied for each of the six spots. As predicted, we found that the majority of worms migrated leftwards towards the magnet. This was apparent because the largest portion of worms was found at the upper left spot (A) while smallest portion was found at the lower right spot (F). Both of these groups were significantly different from chance (point A, *t* = 2.96, p < 0.01; point F, *t* = 4.60, p < 0.001; n=19 assays, Fig 5B). This leftward magnetotaxis trajectory also appears consistent with the independent results by Caldart and Golombek who found that freely moving worms accumulated on the left side of a north-facing up magnet after 15 minutes (Fig 2).

**Figure 5.**
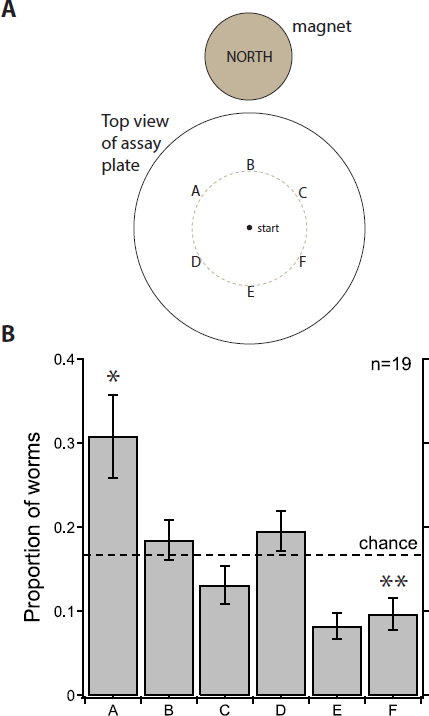
Asymmetric accumulation in six-point magnetotaxis assay. **A)** A group of worms was placed at center of the plate with a north-facing magnet on one side and allowed to move freely. At the end of the 30-minute assay, worms immobilized by azide at six points (A-F) were tallied with the experimenter blind to position of the magnet and identity of the points. **B)** As predicted for a leftward arced trajectory, compared to chance level of accumulation of 0.167, more worms accumulated at point A (p < 0.01) and far fewer worms accumulated at point F (p < 0.001). Bars represent s.e.m. N = 19 assays.

Taken together, this new analysis unifies the migratory patterns observed in all three behavioral assays and yields new predictions strengthening our original findings.

### Magnetic orientation in three-dimensions

In our original study, we observed that different wild *C. elegans* strains isolated from different locations on the earth migrate at a different particular angles relative to magnetic north. For instance, relative to magnetic north, well-fed worms from Britain accumulate on average at 132°, Australian worms at 302°, and Hawaiian worms at 121°. Moreover, when worms were starved, each strain migrated ~180° relative to the preferred angle when well fed. In our study, we made the parsimonious hypothesis that these different angles may relate to the different inclination angle of the earth’s magnetic field at each location; rather than simply following the inclined angle of the earth’s magnetic field, these angles may help worms burrow more directly up or down in 3D rotting vegetation.

Given these results, Landler et al. (2017), and originally Parthasarathy (2015), suggested that if worms simply migrated at a fixed angle relative to the magnetic field, then a population of worms dispersing from a single point outward would form a cone-shaped trajectory when burrowing in three-dimensional space. The apex angle of the cone would 14 be twice the preferred angle and only one line along the cone would aim correctly up or down.

Although we reported that each worm strain migrated at a different *particular angle* relative to the magnetic field when crawling on the 2D agar surface, our hypothesis extrapolating this behavior to three dimensions was admittedly unclear. To clarify, we meant that each worm strain would move in a different *particular* vector relative to the magnetic field. This vector would point along the single vertically aligned line in the cone-shaped trajectory described above. With this behavioral strategy, worms may move optimally in vertical directions.

We were also puzzled how worms migrated at a particular angle to 2D magnetic field in our horizontal plate assay and how they might determine a particular vector relative to a 3D magnetic field. With our new analysis above, however, this unexpected behavior appears to be consistent across all three of our magnetic orientation assays. It is important to clarify that these results provide evidence demonstrating that worms do not simply migrate at a fixed angle with respect to magnetic north. If worms did migrate at a fixed angle to the field in a conical trajectory, then worms would have accumulated at <u>two positions</u> symmetrical about magnetic north on the edge of the horizontal plate. Instead, we found in that worms consistently accumulated at <u>only one position</u>. We observed this result in six out of six conditions – 3 independent wild-type strains in both fed and starved states. Likewise, if worms simply migrated at a fixed angle with respect to magnetic north in the magnetotaxis assay, then they would have moved both towards and away from the magnet in similar proportions based on our new analysis above.

We still do not understand how or why worms behave this way and are not wedded to any particular hypothesis. As the first study of magnetic orientation in *C. elegans*, we do not feel that we have to provide an explanation for something that we do not yet understand. We generally expect that the sensed direction of the magnetic field is combined with allothetic information from an additional sensory cue(s) to disambiguate the cone defined by a single fixed angle with respect to the vector field. For example, *C. elegans* might compute a final 3D vector trajectory by using vector math. The *vector cross product* of two input vectors results in a single output vector. In this case, the worm might use the AFD neurons to sense the magnetic field as a first input vector and allothetic input (e.g. humidity, texture, gaseous and/or thermal gradients) as a second input vector. As long as the two input vectors are not parallel or anti parallel, then their cross product will define a single resultant vector. The worm may also use additional knowledge of its own position or motion (via efference copy or motor signals) to refine a vector calculation. The resultant vector could represent the actual appropriate trajectory vector or some parameter that is used to guide the appropriate trajectory. Examples of nervous systems performing vector math calculations include Schor and Angelaki, 1992; Salinas and Abbott, 1994; and Green and Angelaki, 2004. In short, the worms are sensing something else (or maybe more than one other thing) to pick out the correct single vector with respect to the magnetic field. We acknowledge that this model is speculative, but this is one mathematical mechanism that could be used to distinguish a single trajectory vector. Therefore, the ability of the worms to orient at a specific vector with respect to the magnetic field is categorically possible.

All animals that orient to magnetic fields are believed to combine magnetic cues with additional cues such as light or gravity. We are also currently testing whether additional sensory information including temperature, humidity, or even gravity contributes to this calculation in *C. elegans*. Alternatively, the uniquely complex morphology of the AFD sensory ending and pairing of the two AFD sensory neurons may provide an unexpected mechanism that allows *C. elegans* to determine a specific migratory vector to the field (Clites & Pierce, 2017). The sensory endings of the AFD have an intrinsically rod-shaped character with dozens of nanometer-scale microvilli sprouting in the anterioposterior dimension (Doroquez et al., 2014). This shape gives the neuron the potential to respond by bending or twisting its sensory structure in the presence of a magnetic field if it were found to be associated with iron.

### Physiological evidence for magnetoreception in *C. elegans*

Landler et al., (2017) focused on our behavior experiments without discussing their role in predicting that the AFD sensory neurons responded to magnetic stimuli. After using our behavioral assays to identify the AFD neurons as essential for magnetotaxis and vertical burrowing, we used calcium imaging to show that they respond to magnetic fields with a reliable increase in intracellular calcium (1-2% when worms were physically restrained and 20% when unrestrained) (Vidal-Gadea et al., 2015). We showed that this magnetosensory response did not rely on chemical synapses, but required the presence of a functional tax-4 gene. Only a handful of studies have provided physiological evidence for the nervous system responding to magnetic fields (Pavlova et al., 2011; Wu & Dickman, 2012). To the best of our knowledge, our Vidal-Gadea et al. (2015) study represents the first evidence of a sensory neuron responding to an earth-strength magnetic field. Despite the potential significance of these results to the field of magnetosensation, Landler et al. (2017) did not acknowledge these physiological findings nor critique them.

## Conclusion

Magnetic orientation may be challenging to test in *C. elegans*, but worthwhile to get a foothold in discovering some of the first evidence for cellular and molecular basis for magnetoreception in animals.

## Methods

### Caldart and Golombek Magnetic assay

A modified magnetotaxis assay developed by Caldart and Golombek was performed in Buenos Aires, Argentina. Nematodes were subjected to light:dark cycles (LD, 400:0 lux 12:12 h) under constant temperature (17.5°C). Nematode populations were synchronized to the same developmental stage by the chlorine method (Lewis & Fleming 1995). The harvested eggs were cultured overnight in a 50 ml Erlenmeyer flask with 3.5 ml of M9 buffer (42 mM Na_2_HPO_4_, 22 mM KH_2_PO_4_, 85,5 mM NaCl, 1 mM MgSO_4_) + antibiotic–antimycotic 1× (Gibco, Carlsbad, CA), at 110 rpm, 18.5 °C and under LD 12:12 h conditions.

The next day, L1 larvae were transferred to NGM plates with *E. coli* hb101 to develop up to L4 stage. Then, 500 individuals were transferred to NGM plates with *E. coli* HB101. For the magnetic migration assay, natural magnets (Neodymium) were placed under the plate near the edge, providing an effective magnetic field of 700 Gauss (around 100 times the strength of that of the Earth) on the plate. In order to avoid any magnetic field, the control group was placed into a Faraday cage, and plates were covered with an iron mail with a cape of zinc, of hexagonal design. This mail was covered with aluminum foil for a better magnetic insulation. The experimental plates with the magnet were also covered with aluminum foil, with the magnet inside the package (without contacting the metal).

At the onset of the test, the nematodes were placed in the center of the plates, which were imaged 30 min later with a digital scanner (1800 dpi resolution). The agar plate was digitally divided into 8 octants numbered in a clockwise fashion, (with the magnet in the junction of octants 1 and 8). The number of worms was counted with the particle analyzer plug-in of image-j software, and individuals present in each octant were assessed as the proportion of total worms in the plate. Migration was analyzed with a Rayleigh Test to obtain the clustering as well as the significance of the resulting vector.

Two nematode strains were used in these assays: *C. elegans* strains N2 (Bristol strain, wild-type), and PR671 *tax-2(p691)*, provided by the *Caenorhabditis* Genetics Center.

### Estimation of magnetic field

We used the K&J magnetics magnetic field calculator to approximate field strength over distance and validated the resulting field components with our DC milligauss meter model mgm magnetometor (Alphalab, Utah).

### Preferred migratory direction over time

Worms were assayed for their preferred migratory direction as a function of time in Normal, IL, USA.

#### Animals

30 to 50 day one adult worms were used in each assay as previously described (here and Vidal-Gadea et al., 2015). Animals came from bleach-synchronized plates and were never starved, overpopulated, or infected. Each culture plate contributed to one assay. Worms grew in temperature and humidity controlled room. No experiments were conducted if environmental weather events caused humidity or temperature fluctuations above 5 degrees or % humidity.

#### Assay plate

Animals were transferred using a 0.5-μl droplet of liquid NGM (pH7) into the center of a one day old 10-cm chemotaxis assay plate. Sodium azide was painted with a paintbrush on the circumference of the assay as described before (Vidal-Gadea et al., 2015). Animals were released to behave by carefully soaking the liquid NGM where they were trapped using a small piece of Kimwipe.

#### Magnetic coil system

We used a 1-m^3^ magnetic coil system consisting of three independently powered four-coil Merritt coil system as previously described (Vidal-Gadea et al., 2015) except for one important feature: our new system is double wrapped (Kirschvink, 1992). Temperature and magnetic measurements were performed before and after each experiment to confirm our experimental conditions. A small fan circulated air through the volume of the magnetic coil system to prevent temperature gradients from building up.

#### Magnetic assays

We performed three types of assays. A homogeneous magnetic field of 0.65 Gauss (one earth strength) was produced across the horizontal plate within the test volume (test condition, N = 7). We ran magnetic controls in which the magnetic coil system was used to generate a magnetic field equal and opposite to that of the earth. This cancelled magnetic fields inside the test volume (magnetic control, N = 10). Next we powered our magnetic coil system to generate a 0.65-Gauss field once more but after attaining this field we switched the double-wrapped coil system into its antiparallel configuration where the field generated cancelled itself but produced the same power output as our test condition (current control, N = 6). Before each assay we rotated the magnetic coil system to a random starting position. To determine possible temperature gradients we measured the temperature difference between the center of the assay plates and its edge throughout the assays for each condition as previously reported in Vidal-Gadea et al. (2015).

#### Filming

We used a USB camera (Celestron) driven by Micro-Manager software to film the magnetic assay. Two LED light sources were used to illuminate the filming arena. Test images were obtained and quantified using ImageJ to ensure no brightness gradients were present across the entire filming arena (measuring 40×40mm). Worms were filmed at 1 fps for 100 minutes. Both USB camera and USB lights were wrapped in a grounded faraday fabric made of copper. The same material was used to completely enclose the assay and prevent any electric fields from intruding in our assay.

#### Analysis

Worm centroids were tracked using ImagePro7 object tracking feature by experimenters blind to assay treatment. The heading of each worm was obtained by custom made script in Spike2 (Bainbridge, 2017). We divided the movie into ten windows of equal duration (10 min each). The x and y coordinates of each worm centroid were used to calculate the overall heading of each animal for each time window. We used circular stats toolbox in Matlab (Mathworks) to calculate the average heading for each time window in each assay. These averages from each window were then used to calculate the mean heading of the entire set of assays). For this manuscript, we pooled the headings over 30 minute intervals (time windows 1 through 3, 4-6, 7-9) for a total of 90 minutes out of the 100-minute assays. We used Matlab to obtain the mean heading of the groups of worms over the 30 minute intervals described.

### Six-point magnetotaxis assay

The six-point magnetotaxis assay was performed in Austin, TX, USA and represents a modified version of our previously described magnetotaxis assay (Vidal-Gadea et al., 2015) with two main differences. In the six-point assay, three 1-μL droplets of sodium azide were placed in the upper and lower halves of the plate, rather than one droplet. The droplets (Fig 5A, labeled A-F) were placed 2.5 cm from the center of a 10 cm diameter, agar-filled Petri plate at the following positions (for reference 0° = top center of the plate; 180° = bottom center of the plate): A = 315°; B = 0°; C = 45°; D = 225°; E = 180°; F = 135°. Second, the magnet was moved so that it was adjacent to the plate, rather than directly beneath the upper quadrant of the plate. The magnet was also covered in a 0.5-cm plastic barrier so as to minimize the formation of a temperature gradient across the plate. First day adult worms were washed 3 times in NGM buffer, before being transferred to the center of the assay plate. Excess NGM from the puddle was wicked away, and worms were allowed to migrate for 30 min. At the end of the assay, worms paralyzed at each of the six points were tallied blind to position of the magnet. Significant deviation from chance was assessed using a two-tailed comparison from mean (Zar, 1999).

### Statistics

Vectorial data was analyzed as previously described (Vidal-Gadea et al., 2015) using Circular Toolbox for Matlab (Mathworks). Following Landler et al., (2017), animals were not pooled but each assay was rather treated as a unit. We conducted Rayleigh tests to determine probability of deviation from circular distribution. Non-parametric groups were compared using Mann-Whitney Ranked Sum tests.

## ACKNOWLEDGEMENTS

We wish to acknowledge the *Caenorhabditis* Genetics Center which is supported by the National Institutes of Health, as well as NIH grants to A.V-G. (R15AR068583) and J.P. (R01NS075541 and 1RF1AG057355). D.A.G. and C.S.C. are funded by the National Science Agency, CONICET and University of Quilmes, Argentina.

